# Comparing auditory and visual aspects of multisensory working memory using bimodally matched feature patterns

**DOI:** 10.1101/2023.08.03.551865

**Authors:** Tori Turpin, Işıl Uluç, Parker Kotlarz, Kaisu Lankinen, Fahimeh Mamashli, Jyrki Ahveninen

**Author notes:** equal contribution (Tori Turpin, Işıl Uluç). Corresponding author: Işıl Uluç, Ph.D. CNY 149, 13th St. A.A. Martinos Center for Biomedical Imaging, Department of Radiology, Massachusetts General Hospital, Charlestown, MA 02129.

## Abstract

Working memory (WM) reflects the transient maintenance of information in the absence of external input, which can be attained via multiple senses separately or simultaneously. Pertaining to WM, the prevailing literature suggests the dominance of vision over other sensory systems. However, this imbalance may be stemming from challenges in finding comparable stimuli across modalities. Here, we addressed this problem by using a balanced multisensory retro-cue WM design, which employed combinations of auditory (ripple sounds) and visuospatial (Gabor patches) patterns, adjusted relative to each participant’s discrimination ability. In three separate experiments, the participant was asked to determine whether the (retro-cued) auditory and/or visual items maintained in WM matched or mismatched the subsequent probe stimulus. In Experiment 1, all stimuli were audiovisual, and the probes were either fully mismatching, only partially mismatching, or fully matching the memorized item. Experiment 2 was otherwise same as Experiment 1, but the probes were unimodal. In Experiment 3, the participant was cued to maintain only the auditory or visual aspect of an audiovisual item pair. In two of the three experiments, the participant matching performance was significantly more accurate for the auditory than visual attributes of probes. When the perceptual and task demands are bimodally equated, auditory attributes can be matched to multisensory items in WM at least as accurately as, if not more precisely than, their visual counterparts.

## Introduction

Working memory (WM) plays a crucial role in our cognitive abilities, encompassing the processes of encoding, transiently retaining, and subsequently accessing pertinent external information that guides our goal-oriented behaviors. In the complexities of daily life, inputs accumulate through multiple sensory systems, causing multiple types of information maintained in our WM. Existing theories propose that information is received through distinct modality-specific subsystems of WM (A. Baddeley, 1986; De Renzi & Nichelli, 1975; Quak, London, & Talsma, 2015). Of these subsystems, the visual WM has been considered to have the highest capacity and most precise performance, a characteristic shared by both humans and primates (Bigelow & Poremba, 2014; Colombo & D’Amato, 1986; Hashiya & Kojima, 2001).

A multitude of compelling evidence reinforces the dominance of visual WM. This assertion finds support in numerous studies that have systematically compared various sensory modalities with the visual modality within the realm of human WM. Cohen et al. (2009) examined whether the robust retention capabilities ascribed to visual memory are also found in the auditory domain. Using a range of complex, naturalistic sounds and images with equal memorability, they tested the recognition for sound clips, verbal descriptions, and matching pictures alone. Recognition accuracy in the experiments were consistently and substantially lower for sounds than visual scenes. Similarly, Bigelow and Poremba (Bigelow & Poremba, 2014) provide further corroboration of the visual WM prominence. Their findings indicate that human auditory WM exhibits a notable inferiority in performance when compared to the visual counterpart, especially when dealing with retention periods extending beyond a few seconds. Comparable findings have been reported in behavioral studies in non-human primates (Colombo & D’Amato, 1986; Scott, Mishkin, & Yin, 2012), as well as in recent neurophysiological studies in humans (Wolff, Kandemir, Stokes, & Akyurek, 2020).

However, the marked modality differences in the precision and capacity could represent innate processing differences between auditory and visual WM (Li & Cowan, 2021; Noyce, Cestero, Shinn-Cunningham, & Somers, 2016; Penney, 1989). The traditionally strong emphasis on visual studies might have resulted in designs that favor memoranda in this modality over the others (Lehnert & Zimmer, 2008). For example, in the visual domain, one can encode multiple distinct objects in parallel (Luck & Vogel, 1997). In the auditory domain, WM objects need to be distributed across relatively long periods of time (Bizley & Cohen, 2013), which in itself could increase the memory load (A. D. Baddeley, Thomson, & Buchanan, 1975). Comparisons of performance using classic capacity measures, such as the number of briefly presented items that can be maintained in WM, could thus inherently favor the visual domain.

In the present study, we compared the fidelity of auditory and visual WM by using stimuli that reflect corresponding parameters in each modality, as motivated by previous attempts to directly compare parametric features of these modalities by utilizing simple, manipulable stimuli (Visscher, Kaplan, Kahana, & Sekuler, 2007) (**Fig. 1**). In the visual domain, we chose static Gaussian-windowed sinusoidal gratings*, i.e.*, Gabor patches, a staple stimulus family used to probe visual WM. To guarantee comparable stimulus representations that reflect auditory WM, we utilized frequency-varied broadband sounds, referred to as dynamic ripple sounds, which are resistant to semantic contamination but possess spectrotemporal properties similar to human speech (Depireux, Simon, Klein, & Shamma, 2001). We employed a retrospective cue (retro-cue) paradigm, which helps separate the account of actively maintained WM content from the stimulus history for our primary WM tasks (Backer & Alain, 2012; Kumar et al., 2016; Lepsien & Nobre, 2006; Mamashli et al., 2021; Uluc, Schmidt, Wu, & Blankenburg, 2018). Most importantly, unlike the majority of studies that have utilized similar approaches, in our main experiments, we used a multisensory design to compared the performance of auditory and visual WM to allow modality comparisons across the same trials. We hypothesized that memory for feature information in audition and vision would at least be equally reliable when systematically equated in content and complexity.

**Figure 1.**
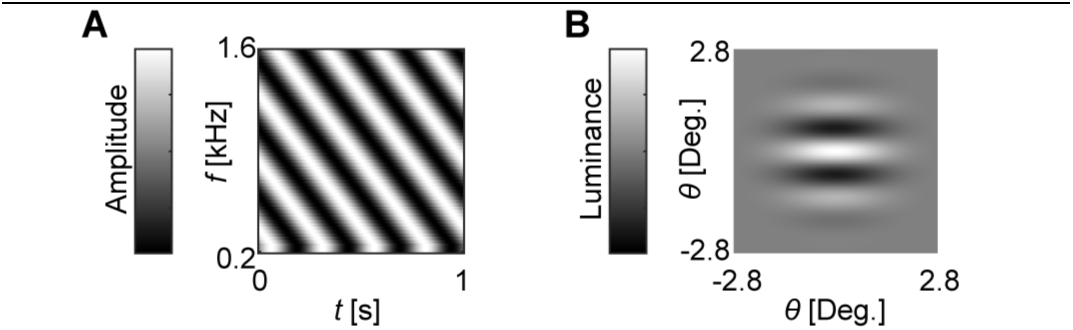
Auditory and visual WM items. (A) An Example for Auditory Stimulus. Dynamic ripple sounds refer to broadband (here, 0.2–1.6 kHz) sounds that are modulated across time (ripple velocity ω; measured in cycles per second) and frequency (spectral density Ω; measured in cycles/octave). (B) An Example for Visual Stimulus. Representations of a Gabor visual gratings plotted as a function of space measured in degrees of visual angle θ around fixation.

## Methods

### General methods for all Experiments

#### Participants

The study protocol was performed in accordance with the Human Research Committee and the Institutional Review Board (IRB) of Massachusetts General Hospital. A total of 40 healthy adult volunteers (age: mean ± standard deviation, SD, 32 ± 12 years; 16 women) participated in at least one of the four Experiments for monetary compensation: 22 subjects took part in Experiment 1 (mean ± SD age 31 ± 11 years; 10 women), 21 subjects in Experiment 2 (mean ± SD age 36 ± 13 years; 7 women), 21 subjects in Experiment 3 (mean ± SD age 36 ± 13 years; 7 women), and 10 subjects (mean ± SD age 34 ± 13 years; 5 women) in the additional Control Experiment. Subjects self-reported no hearing deficits and normal or corrected-to-normal vision. Written informed consent was obtained from each participant prior to experiments.

#### Apparatus

Stimuli were projected from a 24” Dell Model U2410 Monitor with 1920×1200 pixel resolution driven at 59 Hz. Viewing distance at eye-level was 50 cm in front of the subject. The display was affixed at the center of the visual field, subtending a 5.6° visual angle. All visual stimuli were presented in the center of a grey background. Head movements were stabilized with the aid of a chinrest. Sounds were presented binaurally through HD-280 supra-aural headphones (Sennheiser Electronic Corporation). Presentation of the experiment was controlled using Presentation Software (Neurobehavioral Systems, San Francisco, CA).

#### Auditory Stimuli

For all parts of the study, the dynamic ripple waveform sounds were created by layering 20 sinusoids per octave with frequencies ranging from 0.2 kHz to 1.6 kHz. The frequencies, *f*, upon which ripple velocities vary (at a rate of ω cycles per second) are represented by the equation: *s(g, t)* = *D_o_* + *D* ⋅ *cos*[2π(ω*t* + Ω*g*) + ψ] where 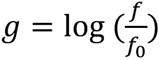, t is time, *D_o_* is baseline intensity, *D* is modulation depth, ω is ripple velocity, Ω is spectral density, and ψ is the phase of the ripple. At presentation, the auditory stimuli featured a fade in and out for 10 ms at the beginning and end of each stimulus.

#### Visual Stimuli

For all parts of the study, the static Gaussian windowed, visual sinusoidal gratings (Gabor patch) stimuli were created by function *g = Acos*(*fz*), where *g* is grating pattern, *A* is maximum contrast of the image, *f* is spatial frequency (cycles per pixel), and *z* is rotated coordinates with angle θ, so that *z = xcosθ + ysinθ*, where *x* and *y* are the coordinates of the image. We used values *A* = 0.2, θ = 90 degrees and image resolution 257×257 (defining the range of *x* and *y*). The grating pattern was masked with a Gaussian function *m = Bexp* – 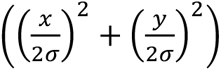, where *B* is a scaling factor and σ is standard deviation. We used values *B* = 1 and σ = 32 pixels (corresponding to 5.6° degrees around fixation). Furthermore, the image was scaled so that the maximum amplitude of the image was 1 and that the edge luminance was exactly 0.5 to blend the patch with the surroundings.

#### Preliminary Detection Thresholds

To ensure that the auditory and visual WM materials were identical in difficulty in all experiments, stimulus-specific preliminary detection tasks controlling for individual baseline differences in sensory discrimination were administered. Subjects received separate tasks in the auditory and visual modalities prior to the memory experiment. Modality task order was counterbalanced according to parity of subject identification number. Just noticeable difference (JND) values were estimated using a 2-down, 1-up staircase algorithm separately in each individual participant. From these values, we generated modality-specific auditory and visual arrays of six stimuli, in which each stimulus was 1.5 JNDs away from its nearest neighbor. All main experiments’ stimuli were drawn from these arrays. The classes 2–5 were used for the WM item stimuli, whereas all classes were used for the probes. Given that stimuli were individually tailored to each participant, baseline stimulus values (stimulus 1 in each array) were constant across the group, whereas subsequent stimulus values 2-6 were adapted for each participant. Three participants whose ripple stimulus classes included stimuli with ripple velocities above 500 cycles/s were excluded from further analysis (Supin et al., 2018).

#### Procedure

Behavioral experiments took place in a quiet, dimly lit room. The experimenter explained instructions prior to administering the task. During explanation, participants had the opportunity to ask for any clarifications they needed. In all experiments, the subjects participated in 10 practice trials and then were left alone to complete the experiment.

### Experiment 1: Bimodal retro-cue task with bimodal probes

22 participants (mean ± SD age 31 ± 11 years; 10 women) of the total cohort took part in Experiment 1. Trials began with a “!” to denote a new sequence. Then, the two sets of concurrently audiovisual stimuli were presented in succession for 1 s each with a 350 ms offset-to-onset interstimulus interval (ISI) in between. This was followed by a 500-ms visual retro-cue (“1” or “2”) that instructed which item to retain in memory over a pseudorandomized subsequent maintenance period (either 2 or 10 s). Then, a probe stimulus was presented for 1 s, at which time participants were to respond to the probe. In the case of non-match probes, the probes’ auditory and/or visual attributes were separated by one step along the feature axis, that is, 1.5 JNDs from the relevant item. The paradigm was split into six sequence blocks, each comprised of 48 trials, adding up to 288 trials in total.

A match denoted audiovisual probe and memory items when they were conceptually perceived as identical entities (same or match trials). Conversely, non-match items were specified by three categories in which; sound features differed (auditory non-match or ANM trials), visual features differed (visual non-match or VNM trials), or both sensory features differed (audiovisual non-match or NM trials). In unimodal non-match trials, the key distinction lay in the dissimilarity of items in terms of their feature representations within the auditory or visual components. ANMs were represented by nonidentical ripple velocities, whereas VNMs were represented by nonidentical Gabor spatial frequencies. Audiovisual NM trials were bimodally nonidentical in stimulus features. Notably, participants were naïve to partial unimodal match conditions as to promote conceptual congruency within the item. An equal number of match and non-match trials (with even distribution between the three non-match conditions) were presented in random order throughout the six sequence blocks. Four stimulus values were drawn from individualized pools of classes among each modality, resulting in four auditory and visual test features culminating in 16 possible audiovisual multifeatured combinations.

Response mapping was counterbalanced according to parity of subject number. Half of the participant pool indicated a “same” response with the right index finger press and a “different” response with the right middle finger press, and the other half of the subjects vice versa. If no response was given within a 1.5 s time window the monitor reminded the participant to respond until a response was received. A feedback was provided by three crosses “+++” to signify a correct response or three dashes “---” for an incorrect response. New trials commenced following an intertrial interval (ITI) of 1 s.

### Experiment 2: Bimodal retro-cue WM task with unimodal probes

21 participants (mean ± SD age 36 ± 13 years; 7 women) of the total cohort took part in Experiment 2. The procedure, instructions, and stimuli were similar to those of Experiment 1, with the only notable difference being that in this experiment, the probe was exclusively either auditory or visual (**Figure 3**). Trials began with a “!” to denote a new sequence. Then, the two sets of concurrently audiovisual stimuli were presented in succession for 1 s each with a 350 ms offset-to-onset interstimulus interval (ISI) in between. This was followed by a 500-ms visual retro-cue (“1” or “2”) that instructed which item to retain in memory over a pseudorandomized subsequent maintenance period (either 2 or 10 s). Then, a probe stimulus, which was either an auditory ripple sound or visual Gabor patch, was presented for 1 s. In the case of non-match probes, the probes’ auditory or visual attributes were separated by one step along the feature axis, that is, 1.5 JNDs from the relevant item. In line with Experiment 1, the experiment comprised of 288 trials that were conducted in 6 sequential, 48-trial blocks.

**Figure 2.**
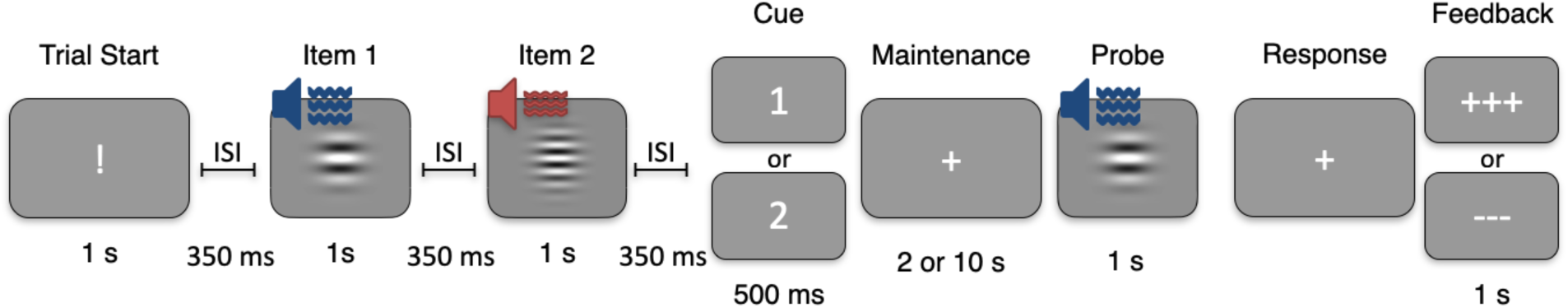
Multimodal Trial timeline. Experiment 1 sequences contained two audiovisual items, presented in a row, with parametrically modulated features. A brief cue indicated the to-be-memorized item (1 or 2), followed by a delay interval of 2 or 10 s. Then, a new bimodal probe was presented. Participants were to determine whether the cued item was the same or different from the probe. The match, ANM, VNM and NM trials as well as 2- and 10-s trials were randomized across the experiment. Retro-cue ‘1’ and ‘2’ trials were also randomized and the retro-cue indicated the to-be-remembered item.

**Figure 3.**
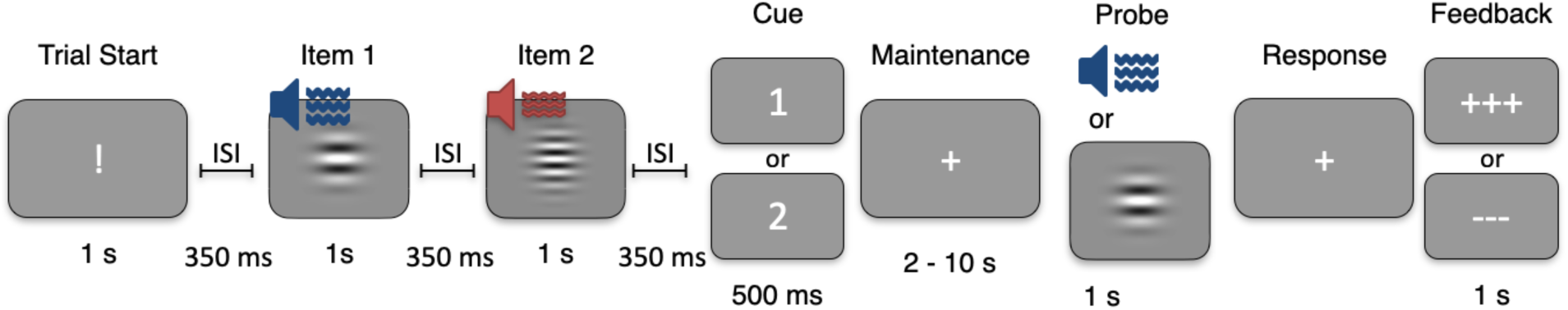
Timeline of the bimodal WM experiment with a unimodal probe. Experiment 2 sequences contained two audiovisual items, presented in a row, with parametrically modulated features. A brief cue indicated the to-be-memorized item (1 or 2), followed by a delay interval of 10 or 2 s. Then, an either auditory or visual unimodal probe was presented. Participants were instructed to determine whether the respective auditory or visual aspect cued item was the same as or different from the probe.

Trials with an auditory probe matching the auditory part of the maintained item were defined as Auditory Match trials and those with a mismatching auditory probe as Auditory Non-Match trials. Trials with a visual probe matching the visual part of the audiovisual memory item were defined as Visual Match trials and those with a mismatching visual probe as Visual Non-Match trials. An equal number of Auditory Match, Auditory Non-Match, Visual Match, and Visual Non-Match were presented in random order throughout the six sequence blocks. The stimulus classes in each modality and the potential combinations of audiovisual stimuli were similar to Experiment 1.

The participants were instructed to press one button if the unimodal probe matched to the respective part of the audiovisual memory item and another button if it did not match the respective part of the audiovisual item. As in Experiment 1, response mapping was counterbalanced according to parity of subject number. Half of the participant pool indicated a “same” response with the right index finger press and a “different” response with the right middle finger press, and the other half of the subjects vice versa. If no response was given within a 1.5 s time window the monitor reminded the participant to respond until a response was received. Feedback was provided by three crosses “+++” to signify a correct response or three dashes “---” for an incorrect response. New trials commenced following an intertrial interval (ITI) of 1 s.

### Experiment 3: Bimodal item, retro-cue to unimodal maintenance, and unimodal probe

21 participants (mean ± SD age 36 ± 13 years; 7 women) of the total cohort took part in Experiment 3. Trials began with a “!” to denote a new sequence (**Figure 4**). Then, one concurrently audiovisual stimulus was presented for 1 s. This was followed by a 500-ms visual retro-cue (“A” or “V”) that instructed which part of the bimodal item to retain in memory over a pseudorandomized subsequent maintenance period (either 2 or 10 s). Subsequently, a probe stimulus was presented, matching the same modality as the to-be-remembered item (i.e., a ripple sound for auditory maintenance and a Gabor patch for visual maintenance). This probe was displayed for a duration of 1 s., and participants were required to provide their response to the probe stimulus at that point. In the case of non-match probes, the probes’ auditory or visual attributes were separated by one step along the feature axis, that is, 1.5 JNDs from the relevant item. Similarly to Experiments 1 and 2, Experiment 3 consisted of 288 trials, which were equally divided into 6 blocks.

**Figure 4.**
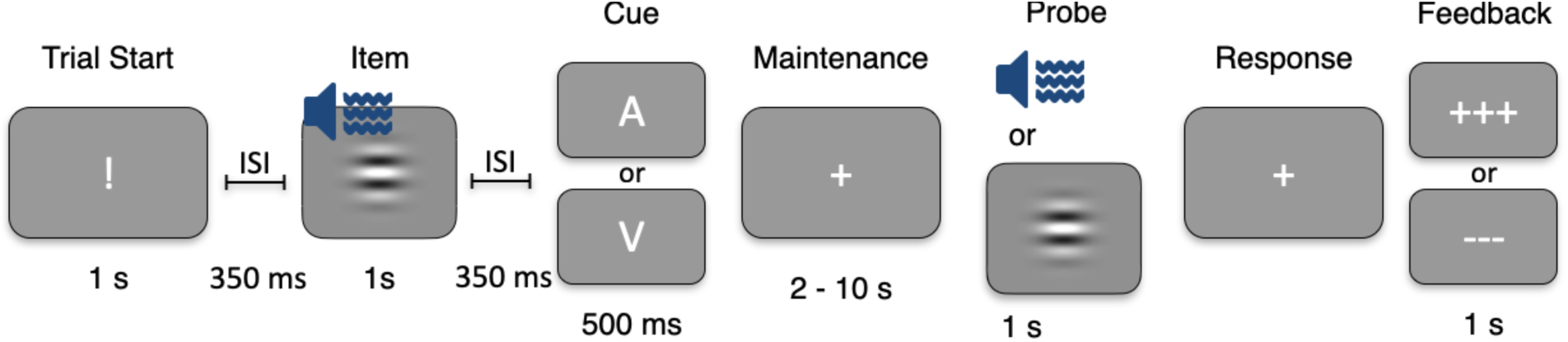
Timeline of the WM experiment with a bimodal item, unimodal maintenance, and unimodal probe. Experiment 3 sequences contained an audiovisual item with parametrically modulated features. A brief cue indicated the to-be-memorized modality (auditory or visual), followed by a 2 s- or 10 s-delay interval. Then, a probe of the cued modality was presented.

Trials with an auditory probe matching the maintained auditory item were defined as Auditory Match trials and those with a mismatching auditory probe as Auditory Non-Match trials. Trials with a visual probe matching the visual memory item were defined as Visual Match trials and those with a mismatching visual probe as Visual Non-Match trials. An equal number of Auditory Match, Auditory Non-Match, Visual Match, and Visual Non-Match were presented in random order throughout the six sequence blocks. Four stimulus values were determined from individualized pools of classes within each modality. This selection yielded four unique auditory and visual test features, ultimately giving rise to a total of 16 potential combinations of audiovisual multifeatured stimuli.

The participants were instructed to press one button if the unimodal probe matched to unimodal aspect of the memory item and another if it did not match that aspect. Response mapping was counterbalanced according to parity of subject number. Half of the participant pool indicated a “same” response with the right index finger press and a “different” response with the right middle finger press, and the other half of the subjects vice versa. If no response was given within a 1.5 s time window the monitor reminded the participant to respond until a response was received. Feedback was provided by three crosses “+++” to signify a correct response or three dashes “---” for an incorrect response. New trials commenced following an intertrial interval (ITI) of 1 s.

### Control Experiment

As a control, 10 participants (mean ± SD age 34 ± 13 years; 5 women) took part in separate unimodal WM experiments for auditory and visual sensory modalities (**Figure 5**). Each trial began with two unimodal sample stimuli (i.e., ripple sounds for auditory experiment and Gabor patches for visual experiment) in a row, which was followed by a cue to denote the relevant stimulus according to serial position. A randomized short or long retention interval ensued, after which a probe was either identical (Match trial) or different (Non-Match trial) to the relevant, cued stimulus. The order in which task types were administered (auditory first, visual second; or visual first, auditory second) was counterbalanced across participants. The unimodal retro-cue task featured 24 trials across six blocks amounting to 144 trials in total.

**Figure 5.**
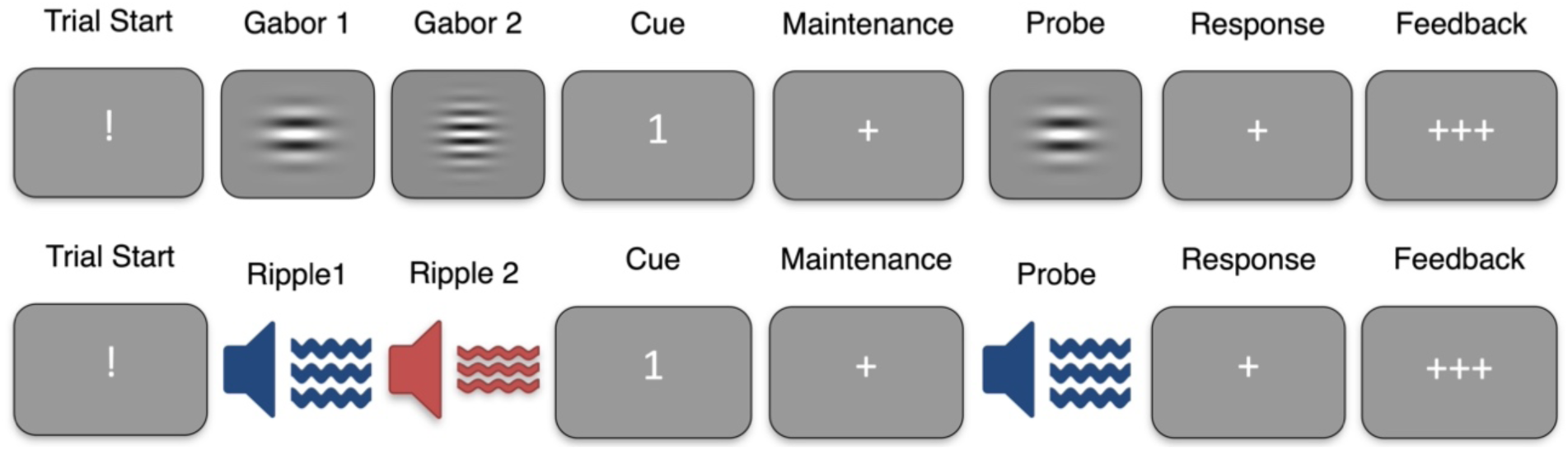
Unimodal Trial timelines, Control Experiment. Visual and auditory presentations in unimodal tasks are displayed in the first and second rows, respectively. Visual trials tested memory for visual Gabor patches while auditory trials tested memory for ripple sounds. Experiment timing and stimulus durations were matched exactly to the three main experiments.

### Quantification of behavioral performance and statistical analysis

In all analyses, the dependent variable was behavioral sensitivity or d’ quantified based on the signal detection theory (Stanislaw & Todorov, 1999) using the hit rate (HR; proportion of “yes” responses for matching Probes) and false alarm rates (FAR; proportion of “yes” responses for non-matching Probes). All behavioral and statistical analyses were conducted in MATLAB using built-in functions and custom code (Trujillo-Ortiz, Hernandez-Walls, & Trujillo-Perez, 2004, 2006). To deall with multiple comparison problems, all reported p-values were corrected using the false discorvery rate (FDR) procedure of Benjamini and Hochberg (Benjamini & Hochberg, 1995; Groppe, 2017).

*Experiment 1* had a three-way within-subject design, which compared the differences in d’ values across these three Conditions (NM, ANM, or VNM), while controlling for the effects of Cued Item (item 1 or 2) and the Maintenance Duration (2 or 10 s). For each Cued Item and Maintenance Duration condition, the d’ values were calculated separately using the FAR values obtained from ANM, VNM, and NM trials using the formula *d’* = z(*HR*) - z(*FAR*_Condition_), where z refers to the inverse of the normal cumulative distribution function. The resulting values were then entered in to a three-way repeated-measures ANOVA testing for the main effects of Condition, Cued Item, and Maintenance Duraction as well as their interactions. More detailed comparisons were also calculated to determine the differences between ANM and VNM conditions across the delay conditions. To correct for multiple comparisons, we adjusted all *p* values assessing the FDR (Benjamini & Hochberg, 1995).

*Experiment 2* had a three-way within-subject design, which compared the differences in d’ values across these two Conditions (Auditory or Visual probe), while controlling for the effects of Cued Item (item 1 or 2) and the Maintenance Duration (2 or 10 s). For each Cued Item and Maintenance Duration condition, the d’ values were calculated separately using the HR and FAR values obtained from auditory or visual trials using the formula *d’* = z(*HR_Condition_*) - z(*FAR*_Condition_), where z refers to the inverse of the normal cumulative distribution function. The resulting values were then entered in to a three-way repeated-measures ANOVA testing for the main effects of Condition, Cued Item, and Maintenance Duration as well as their interactions. To correct for multiple comparisons, we adjusted all *p* values assessing the false discovery rate (FDR) (Benjamini & Hochberg, 1995).

*Experiment 3* involved a two-way within-subject design, which compared the differences in d’ values across two Conditions (Auditory or Visual, cued and probed unimodally), while controlling for the effect of Maintenance Duration (2 or 10 s). For each Maintenance Duration condition, the d’ values were calculated separately using the HR and FAR values obtained from auditory or visual trials using the formula *d’* = z(*HR_Condition_*) - z(*FAR*_Condition_), where z refers to the inverse of the normal cumulative distribution function. The resulting values were then entered in to a two-way repeated-measures ANOVA testing for the main effects of Condition and Maintenance Duraction as well as their interactions. For each Maintenance Duration, the differences across the two conditions were analyzed using paired *t*-tests. To correct for multiple comparisons, we adjusted all resulting ANOVA and *t*-test *p* values assessing the false discovery rate (FDR) (Benjamini & Hochberg, 1995).

*The Control Experiment* involved a three-way within-subject design, which compared the differences in d’ values across these two Conditions (Auditory or Visual), while controlling for the effects of Cued Item (item 1 or 2) and the Maintenance Duration (2 or 10 s). For each Cued Item and Maintenance Duration condition, the d’ values were calculated separately using the HR and FAR values obtained from auditory or visual trials using the formula *d’* = z(*HR_Condition_*) - z(*FAR*_Condition_), where z refers to the inverse of the normal cumulative distribution function. The resulting values were then entered in to a three-way repeated-measures ANOVA testing for the main effects of Condition, Cued Item, and Maintenance Duraction as well as their interactions. To correct for multiple comparisons, we adjusted all *p* values assessing the false discovery rate (FDR) (Benjamini & Hochberg, 1995).

## Results

### Experiment 1: Bimodal retro-cue task with bimodal probes

A group of 22 healthy neurotypical participants performed a fully bimodal reto-cue working memory task with multisensory auditory ripple velocity and visual spatial frequency content. The main finding in Experiment 1 was the participants’ higher sensitivity to reject auditory (ANM) than visual (VNM) non-match probes, which mismatched the audiovisual item in only one sensory domain.

**Figure 4** shows the means and standard errors of means (SEM) of d’ values calculated by using the FAR of either NM, ANM, or VNM trials, separately for the two Cued Item trial types (Cued to 1st, Cued to 2nd) and the two Maintenance Durations (2 s or 10 s). According our three-way repeated-measures ANOVA, there were significant main effects for Condition (*F*_2,42_ = 28.2, *p*_FDR_<0.001) and for Maintenance Duration (*F*_1,21_ = 33.9, *p*_FDR_<0.001), as well as a significant interaction between these two factors (*F*_2,42_ = 4.6, *p*_FDR_<0.05). *Most importantly*, the subsequent ANOVA planned comparisons suggested that the d’ values were significantly larger in the ANM than in the VNM Condition both for both the shorter 2-s (*F*_1,21_ = 24.8, *p*_FDR_<0.001) and the longer Maintenance Durations (*F*_1,21_ = 11.7, *p*_FDR_<0.01). These results suggest a significant difference in the participants ability to detect mismatches in ANM and VNM probes, irrespective of any other intervening effects, including the Cued item and Maintenance Duration.

As depicted in **Figure 4**, the group-average d’ difference between trials Cued to the first vs. second item appeared to be more prominent when the Maintenance Duration was shorter (2 s) than when it was longer (10 s). Indeed, whereas the main effect of Cued Item was non-significant in our main three-way ANOVA, a significant interaction occurred between the Cued Item and Maintenance Duration (*F*_1,21_ = 9.5, *p*_FDR_<0.05). However, the two-way interaction between the Cued Item and Condition, as well as the three-way interaction between Condition, Cued Item, and Maintenance Duration, remained non-significant. In other words, any effects related to Condition (NM, ANM, VNM) did not seem to stem from differences related to whether the participant was cued to maintain the Item 1 or 2.

**Table 1** details the group averages and standard deviations (SD) of d’ values in Experiment 1. **Table 2** shows the group averages and SDs of the respective HR (Match trials) and FAR (Non-Match trials) values.

**Figure 4.**
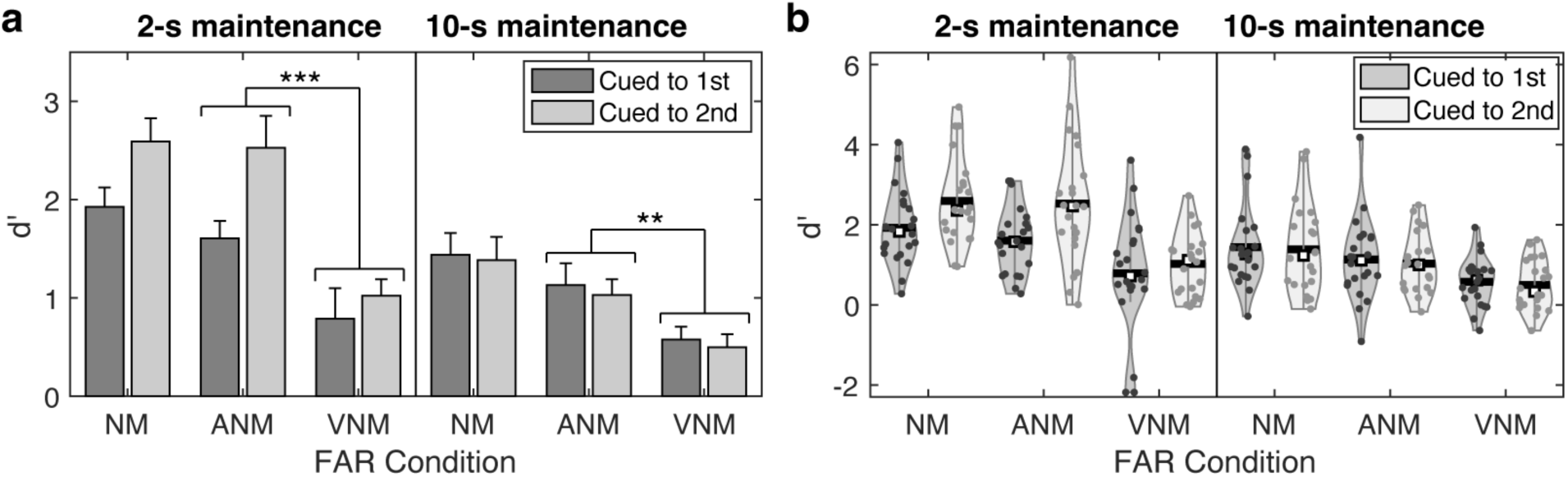
Sensitivity values (d’) in the fully bimodal retro-cue Experiment 1. (**a**) Group mean and SEMs (error bars) of the sensitivity values d’. According to our repeated-measures ANOVAs, the d’ values in the significantly smaller for d’ values calculated using the FAR from VNM than ANM condition in both trials with 2-s and 10-s maintenance period, irrespective of whether the participant was cued to maintain the first or second item pair. More detailed statistical results are provided in the main text (*** *p*_FDR_<0.001) and (** *p*_FDR_<0.01). (**b**) Violin plots of the distributions of d’ values in each condition. The dots are individual participants’ observations. The white squares mark the group medians. The horizontal bars refer to the group means.

**Table 1.** Group mean ± SD values of d’ values in Experiment 1.

| Maintenance | Cued to | NM |  | ANM |  | VNM |  |
| --- | --- | --- | --- | --- | --- | --- | --- |
| 2 s | 1<sup>st</sup> | 1.93 | ± 0.93 | 1.61 | ± 0.83 | 0.79 | ± 1.46 |
| 2 s | 2<sup>nd</sup> | 2.59 | ± 1.1 | 2.53 | ± 1.53 | 1.02 | ± 0.79 |
| 10 s | 1<sup>st</sup> | 1.44 | ± 1.04 | 1.13 | ± 1.03 | 0.58 | ± 0.61 |
| 10 s | 2<sup>nd</sup> | 1.39 | ± 1.1 | 1.03 | ± 0.74 | 0.5 | ± 0.63 |

**Table 2.** Group mean ± SD of HR and FAR values in Experiment 1.

| *Maintenance* | *Cued to* | *HR* |  | *FAR<sub>NM</sub>* |  | *FAR<sub>ANM</sub>* |  | *FAR<sub>VNM</sub>* |  |
| --- | --- | --- | --- | --- | --- | --- | --- | --- | --- |
| 2 s | 1<sup>st</sup> | 0.9 | ± 0.08 | 0.34 | ± 0.17 | 0.45 | ± 0.19 | 0.66 | ± 0.22 |
| 2 s | 2<sup>nd</sup> | 0.9 | ± 0.12 | 0.21 | ± 0.14 | 0.25 | ± 0.23 | 0.66 | ± 0.19 |
| 10 s | 1<sup>st</sup> | 0.69 | ± 0.13 | 0.26 | ± 0.19 | 0.32 | ± 0.2 | 0.49 | ± 0.21 |
| 10 s | 2<sup>nd</sup> | 0.64 | ± 0.19 | 0.23 | ± 0.14 | 0.3 | ± 0.15 | 0.48 | ± 0.24 |

### Experiment 2: Bimodal retro-cue WM task with unimodal probes

In Experiment 2, a group of 21 participants performed a bimodal retro-cue WM task with similar bimodal item pairs and retro-cues to those in Experiment 1, but with unimodally auditory or visual probes. In this experiment, which did not have bimodal competition at the probe matching stage, we found no significant d’ differences between the auditory and visual probe conditions.

**Figure 5** shows the means and SEMs of d’ values calculated by for auditory and visual probe trials, separately for the two Cued Item types (Cued to 1st, Cued to 2nd) and the two Maintenance Durations (2 s or 10 s). In our three-way (Condition x Cued Item x Maintenance Duration) repeated-measures ANOVA, all main effects and interactions involving the Condition (auditory vs. visual probe trials) were non-significant. The only significant effects were the main effect of Cued Item (*F*_1,20_ = 10.5, *p*_FDR_<0.05) and the main effect of Maintenance Duration (*F*_1,20_ = 32.7, *p*_FDR_<0.001).

**Table 3** lists the group averages and standard deviations (SD) of d’ values in Experiment 2. **Table 4** shows the group averages and SDs of the respective HR (Match trials) and FAR (Non-Match trials) values.

**Figure 5.**
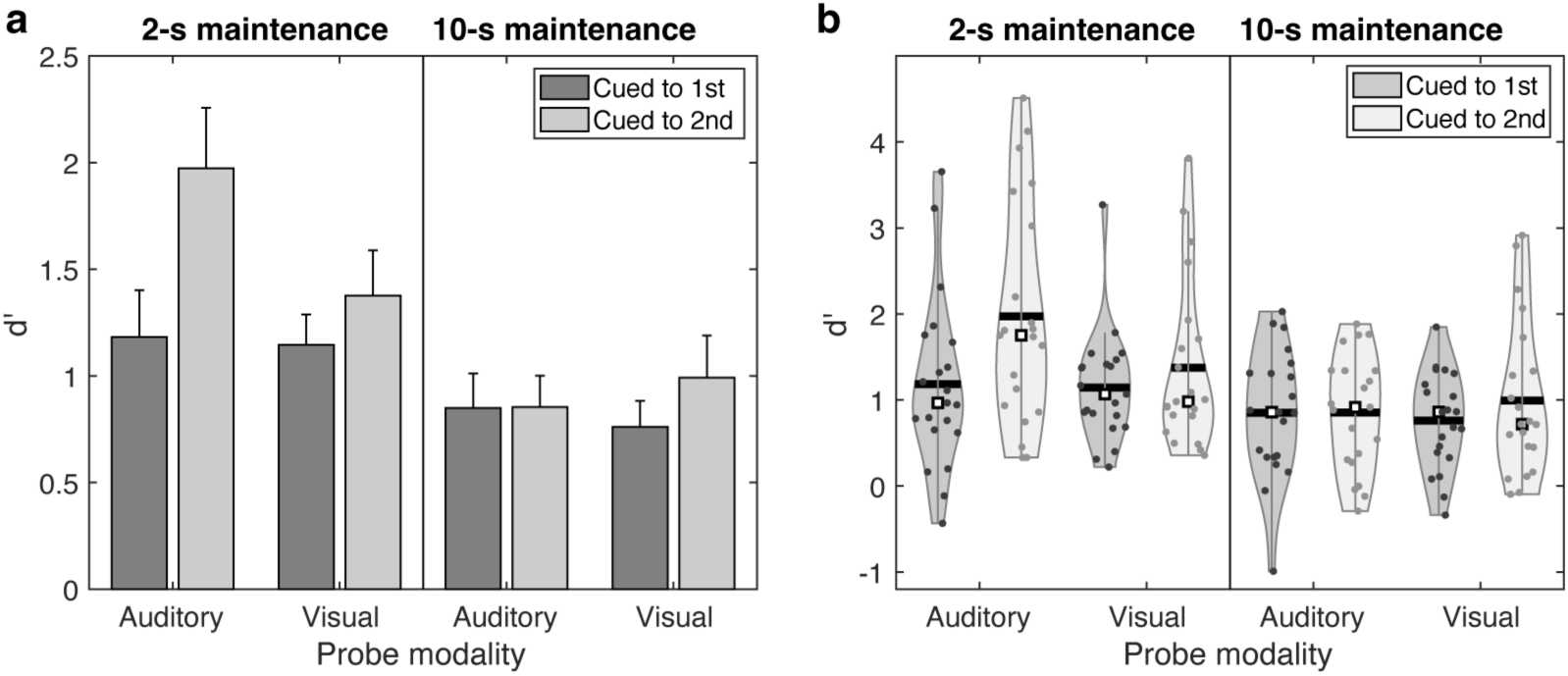
Sensitivity values (d’) in the retro-cue WM Experiment 2, with bimodal items and unimodal A or V probes. (**a**) Group mean and standard errors of the mean (SEM; error bars) of the sensitivity values d’. (**b**) Violin plots of the distributions of d’ values in each condition. The dots are individual participants’ observations. The white squares mark the group medians. The horizontal bars refer to the group means.

**Table 3.** Group mean ± SD values of d’ values in trials with auditory and visual probes in Experiment 3.

| *Maintenance* | *Cued to* | *Auditory* |  | *Visual* |  |
| --- | --- | --- | --- | --- | --- |
| 2 s | 1<sup>st</sup> | 1.18 | ± 1 | 1.15 | ± 0.65 |
| 2 s | 2<sup>nd</sup> | 1.97 | ± 1.29 | 1.38 | ± 0.98 |
| 10 s | 1<sup>st</sup> | 0.85 | ± 0.74 | 0.76 | ± 0.56 |
| 10 s | 2<sup>nd</sup> | 0.85 | ± 0.68 | 0.99 | ± 0.9 |

**Table 4.**
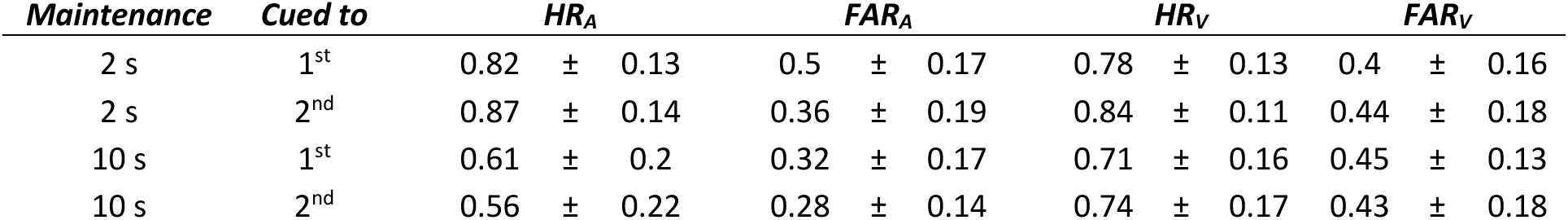
Group mean ± SD values of HR and FAR in trials with auditory and visual probes in Experiment 3.

### Experiment 3: Bimodal item, retro-cue to unimodal maintenance, and unimodal probe

A group of 21 participants performed a WM task where they were first presented with a bimodal ripple sound/Gabor patch pattern, then retro-cued to maintain either the auditory (A) or visual (V) aspect of this bimodal stimulus. Finally, they received a respective unimodal A or V probe to be determined if it is same or different to the memorized aspect (Experiment 3). Figure 6 shows the group averages and SEMs of d’ values in the different task conditions in Experiment 3. According to our two-way repeated-measures ANOVA, there was a statistically significant advantage for A vs. V performance in Experiment 3, as suggested by a significant main effect of Condition (*F*_1,20_ = 17.7, *p*_FDR_<0.001). However, the ANOVA also suggested a significant interaction between the Condition and Maintenance Duration (*F*_1,20_ = 6.4, *p*_FDR_<0.05), suggesting that the difference between A and V conditions changed as a function of the maintenance period. We therefore conducted two additional comparisons of d’ values within each delay condition, which suggested that A vs. V difference was significant in trials with the shorter (*t*_20_=5.2, *p*_FDR_<0.001) maintenance period but remained as a statistically non-significant trend in those with the longer maintenance (*t*_20_=1.8, *p*_FDR_=0.08). Finally, consistent with the other Experiments, there was a significant ANOVA main effect of Maitenance Duration (*F*_1,20_ = 38.3, *p*_FDR_<0.001).

**Figure 6.**
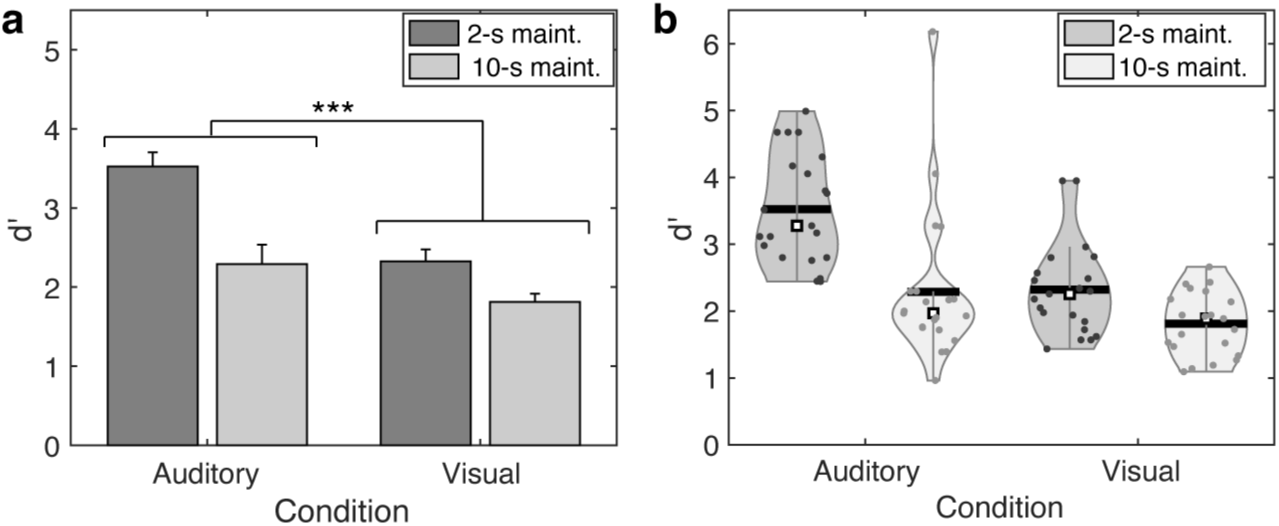
Sensitivity values (d’) in the retro-cue WM Experiment 3, which had bimodal items, retro-cues to unimodal A or V maintenance, and respective unimodal A or V probes. (**a**) Group mean and standard errors of the mean (SEM; error bars) of the sensitivity values d’. According to the ANOVA main effect, the group-average d’ values were significantly smaller for V than A trials. (**b**) Violin plots of the distributions of d’ values in each condition. The dots are individual participants’ observations. The white squares mark the group medians. The horizontal bars refer to the group means. (*** *p*_FDR_ < 0.05)

**Table 5** shows the group averages and standard deviations (SD) of d’ values in Experiment 3. The group averages and SDs of the respective HR (Match trials) and FAR (Non-Match trials) values can be found in **Table 6**.

**Table 5.** Group mean ± SD values of d’ values in auditory and visual conditions in Experiment 3.

| *Maintenance* | *Auditory* |  | *Visual* |  |
| --- | --- | --- | --- | --- |
| 2 s | 3.52 | ± 0.82 | 2.29 | ± 1.13 |
| 10 s | 2.32 | ± 0.69 | 1.81 | ± 0.47 |

**Table 6.** Group mean ± SD values of HR and FAR in auditory and visual conditions in Experiment 3.

| *Maintenance* | *HR<sub>A</sub>* |  | *FAR<sub>A</sub>* |  | *HR<sub>V</sub>* |  | *FAR<sub>V</sub>* |  |
| --- | --- | --- | --- | --- | --- | --- | --- | --- |
| 2 s | 0.96 | ± 0.04 | 0.12 | ± 0.09 | 0.91 | ± 0.06 | 0.22 | ± 0.09 |
| 10 s | 0.76 | ± 0.15 | 0.11 | ± 0.08 | 0.82 | ± 0.08 | 0.22 | ± 0.11 |

### Control Experiment

As a control analysis, we employed the retro-cue procedure independently for the auditory and visual modalities. This enabled us to test direct comparisons of unimodal working memory performance across a cohort of 10 participants (Figure 7). According to our three-way (Condition by Cued item by Maintenance Duration) repeated-measures ANOVA, the main effect of WM modality (A vs. V) on the behavioral d’ was clearly non-significant (*F*_1,9_ = 0.1, *p*_FDR_=0.79). There was a significant interaction between the Condition and Maintenance Duration (*F*_1,9_ = 22.7, *p*_FDR_<0.01). However, according to more detailed planned comparisons, the differences of d’ values between the A and V modalities were non-significant within both the 2-s (*F*_1,9_ = 4.0, *p*_FDR_<0.19) and 10-s (*F*_1,9_ = 3.0, *p*_FDR_<0.19) delay conditions. (Notably, these modality-comparisons between the means within each delay condition would have remained non-significant also without the post-hoc correction conducted using the FDR approach.) Consistent with all other Experiments, there was a significant main effect of Maintenance Duration (*F*_1,9_ = 34.8, *p*_FDR_<0.01). All other ANOVA main effects and interactions remained non-significant. The group means and SDs of the d’ values s are shown in **Table 7** and the respective HR and FAR values in **Table 8**.

**Figure 7.**
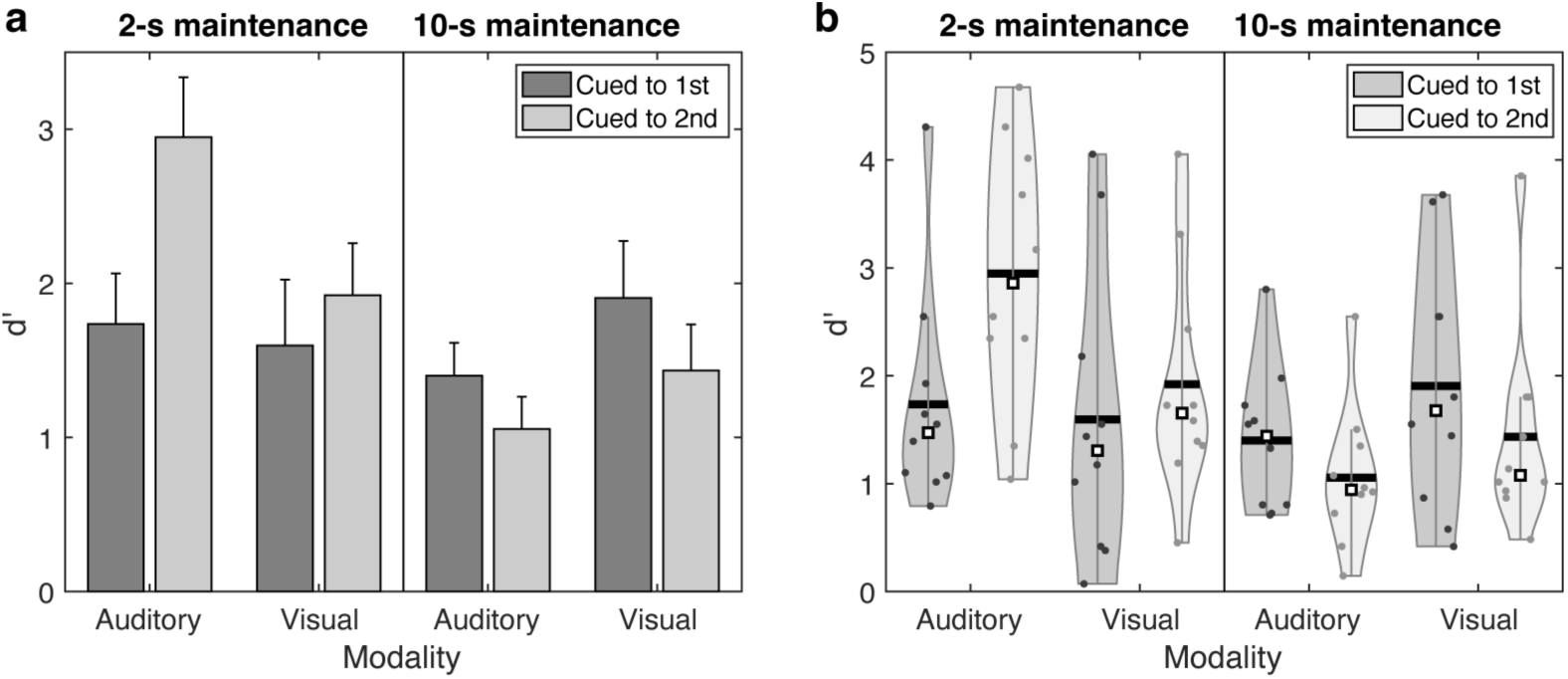
Sensitivity values (d’) in the fully unimodal Control Experiment. (**a**) Group mean and standard errors of the mean (SEM; error bars) of the sensitivity values d’. No significant effects of WM modality (A vs. V) were found in the repated-measures ANOVA. (**b**) Violinplots of the distributions of d’ values in each condition. The dots are individual participants’ observations. The white squares mark the group medians. The horizontal bars refer to the group means.

**Table 7.** Group mean ± SD values of d’ values in auditory and visual conditions in Control Experiment.

| *Maintenance* | *Cued to* | *Auditory* |  |  | *Visual* |  |  |
| --- | --- | --- | --- | --- | --- | --- | --- |
| 2 s | 1st | 1.74 | ± | 1.04 | 1.6 | ± | 1.35 |
| 2 s | 2nd | 2.95 | ± | 1.23 | 1.92 | ± | 1.07 |
| 10 s | 1st | 1.4 | ± | 0.68 | 1.91 | ± | 1.17 |
| 10 s | 2nd | 1.06 | ± | 0.66 | 1.43 | ± | 0.94 |

**Table 8.** Group mean ± SD values of HR and FAR values in auditory and visual conditions in Control Experiment.

| *Maintenance* | *Cued to* | *HR<sub>A</sub>* |  |  | *FAR<sub>A</sub>* |  |  | *HR<sub>V</sub>* |  |  | *FAR<sub>V</sub>* |  |  |
| --- | --- | --- | --- | --- | --- | --- | --- | --- | --- | --- | --- | --- | --- |
| 2 s | 1<sup>st</sup> | 0.81 | ± | 0.14 | 0.27 | ± | 0.13 | 0.79 | ± | 0.16 | 0.34 | ± | 0.13 |
| 2 s | 2<sup>nd</sup> | 0.89 | ± | 0.1 | 0.16 | ± | 0.17 | 0.83 | ± | 0.12 | 0.29 | ± | 0.2 |
| 10 s | 1<sup>st</sup> | 0.64 | ± | 0.17 | 0.18 | ± | 0.11 | 0.74 | ± | 0.19 | 0.22 | ± | 0.17 |
| 10 s | 2<sup>nd</sup> | 0.72 | ± | 0.18 | 0.35 | ± | 0.12 | 0.8 | ± | 0.13 | 0.36 | ± | 0.19 |

## Discussion

We studied the sensory dominance in WM between auditory and visual modalities in a series of experiments that were designed to control for the complexity and difficulty in memory items across modalities. To this aim, we examined the participants sensitivity d’ (Green & Swets, 1966; Stanislaw & Todorov, 1999) to detect matches/mismatches between maintained audiovisual, auditory, and/or visual WM items and subsequent probes in three different bimodal retro-cue WM tasks and in one unimodal control experiment. The materials consisted of non-conceptual auditory and visual feature patterns that were matched for behavioral discrimination difficulty. In the three main experiments, we aimed to disentangle the differences in auditory and visual WM for the perceptual and WM encoding as well as retrieval. Experiment 1 involved bimodal competition at all processing stages. Experiment 2 had the same encoding and maintenance demands as Experiment 1, but the probes were (pseudo-randomly) either auditory or visual alone, thus removing the bimodal competition from the retrieval/decision stage. In Experiment 3, the participants were cued to maintain and match materials in only one of the two modalities. The main finding was that under bimodal competition, the participants were significantly more sensitive to detect auditory than visual mismatches to bimodal probes, when only one of the sensory domains was non-matching to the WM item (Experiment 1). This effect was not present in Experiment 2 that was otherwise similar but lacked the bimodal competition at the probe matching stage. A statistically significant auditory benefit was also observed in Experiment 3, which involved unimodal maintenance and probe matching, suggesting an auditory benefit also in WM maintenance, which was most prominent at the shorter 2-s maintenance durations.

The retro-cue audiovisual paradigm of Experiment 1 required the participants to encode bi-sensory information into WM, actively maintain two intermodally competing memories, and then retrieve the relevant WM features for comparison with a bimodal probe stimulus. For this task, in addition to memory maintenance, adjacent cognitive processes such as divided attention and change detection are intrinsically recruited and utilized during these processing stages (Cowan, 1988; Lavie, 2005). Hence, one explanation for the subjects’ better sensitivity in auditory than visual change in probes could thus be attributed to the multisensory load of maintaining multiple sensory items at one point in time. Given the limited capacity nature of WM, it is possible that intersensory competition for limited storage elicited cross-modal interference effects and led to disadvantage with visual items, which have been reported to be susceptible to interruption in memory and attention (Li & Cowan, 2021).

In a similar vein, the results of classic studies, which compared the behavioral effects of attention cues, suggest that in a multimodal design auditory stimuli tend to be more alerting than visual stimuli (Harvey, 1980). It is thus possible that the present auditorily non-matching (but visually matching) ANM probes resulted in stronger bottom-up activation of the decision-making circuits needed for responding to the retrieval task than their VNM counterparts (Harvey, 1980; Posner, Nissen, & Klein, 1976; Turatto, Benso, Galfano, & Umilta, 2002). This line of thinking can be supported by recent studies, which suggest that auditory change detection is less reliant on sustained selective attention than visual change detection (Demany, Semal, Cazalets, & Pressnitzer, 2010). Such an auditory superiority of bottom-up type change detection has been attributed to an implicit (sensory) memory buffer, which might be considerably weaker in the visual domain (Demany, et al., 2010). However, the present retro-cueing task is believed to control for effects explainable by sensory buffers alone (Lepsien & Nobre, 2006). Further, the potential modality differences in sensory buffers were statistically controlled in all our analyses, by using the Cued Item factor in our statistical models. Nevertheless, modality differences in attention-independent change detection systems could have facilitated sensitivity to auditory change in probes under inter-sensory competition, with the focus of selective attention being shifted back and forth between auditory and visual WM representations.

The results of Experiment 2, where the memorized items were bimodal but the probe was unimodal, suggest a similar interpretation although with some reservations. That is, we found no significant difference in d’ in the case of unimodal probes when there is a bimodal competition during maintenance. These results by themselves would lead us to think that the auditory dominance stems from a sensory competition in retrieval (Harvey, 1980; Robinson et al, 2018). However, a significant auditory benefit was also detected in Experiment 3, which involved a bimodal item presentation but a unimodal maintenance. Although these results do not rule out intersensory competition in the encoding period, the unimodal retrieval in Experiment 3 suggests that sensitivity to auditory probes might be better than the sensitivity to visual ones in bimodal competitions cases. It should, however, be noted that the auditory benefit in Experiment 3 was more prominent in the shorter, 2-s maintenance trials than in the 10-s maintenance trials.

A culprit for differences in WM performance for different sensory modalities could reflect methodological challenges in matching the relative difficulty of auditory vs. visual items and processes (Ward, Avons, & Melling, 2005). To compare the different modalities on an equal footing, the auditory and visual stimulus sets were homogenized in discriminability by using comparable discrimination threshold tests (Mamashli, et al., 2021; Visscher, et al., 2007). Here, the matching of intermodal difficulty was enabled via a threshold test that was similar for auditory and visual stimuli. As discussed in previous studies from which the present stimuli were adapted, such JND estimates could still have become easier for the auditory than visual stimuli (Visscher, et al., 2007). Such a bias could follow from that the participants are, inherently, more familiar in comparing stimuli in one modality vs. another. Hence after the first step of calculating the JNDs for each modality separately, separate unimodal WM Control Experiments on same subjects for both modalities were conducted. In the unimodal comparisons, no significant differences were found in the d’ in auditory vs visual WM performances. We observed a performance drop in both visual and auditory from 2 s condition to 10 s condition but within those conditions or overall, there were no significant modality differences. This result is, in principle, consistent with previous intermodal comparisons using unimodal designs, which suggest that with matched feature content and difficulty, performance of auditory and visual WM follow relatively similar functional principles (Visscher, et al., 2007; Ward, et al., 2005).

There could be modality differences in sensory buffers, which precede active WM processing (Bradley & Pearson, 2012; Näätänen, 1992; Sakitt, 1976; Sperling, 1960; Watkins & Watkins, 1980). For example, the popular belief is that the auditory “echoic” memory (at least 4 s; see, e.g., Watkins & Watkins, 1980) is more persistent than its visual “iconic” counterpart (1 s or less; see, e.g., Bradley & Pearson, 2012). In the present study, we examined this potential bias by analyzing the d’ values separately for trials where the participant was cued to maintain the first vs. second item of the retro cue paradigm. Indeed, there was some evidence that difference between Cued Item 1 and Cued item 2 trials was somewhat more prominent in the auditory than visual domain. However, the difference between Cued Item 1 and 2 trials vanished in both modalities with the longer 10-s maintenance period. Most importantly, using Cued Item as a nuisance factor in our Experiment 1 did not change the main result, suggesting a benefit for detecting auditory vs. visual mismatches under bimodal competition.

A recent modality comparison between auditory, visual, and tactile WM found that the modality differences depend on the maintenance duration, with the auditory WM becoming progressively weaker than the two other modalities at delays of 8 s or longer (Bigelow & Poremba, 2014). In the present study, significant interactions between the modality and maintenance duration were found in the two last experiments that had unimodal maintenance, Experiment 3 and Control Experiment. However, in Experiment 3, which involved a bimodal item presentation and unimodal WM maintenance and retrieval, the trend was still toward an auditory benefit even in the longer 10-s maintenance condition. On the other hand, despite this interaction, no significant differences between the auditory and visual sensitivities were observed in the unimodal Control Experiment. Most importantly, the main result, auditory benefit in detecting mismatches in bimodally incongruent WM probes to audiovisual WM items, was clearly significant even at the longer 10 s delay. Further studies are, however, necessary to determine how these auditory benefits might evolve at longer than 10 s maintenance durations.

## Conclusion

Our results demonstrate that auditory attributes of multisensory WM items can be retrieved as well as if not more precisely than their visual counterparts during inter-sensory competition. No modality differences were observed between auditory and visual WM performance in our unimodal control task. These results could provide a new perspective to the ongoing debate on modality differences in human WM performance.

## Acknowledgments

This work was supported by the NIH grants R01DC016915, R01DC016765, R01DC017991, and P41EB015896.

## Notes

### Competing Interest Statement

The authors have declared no competing interest.

### Summary of Updates

The author added in the last submission is not added into the online version

